# An increase in dendritic plateau potentials is associated with experience-dependent cortical map reorganization

**DOI:** 10.1101/2020.07.13.200923

**Authors:** Stéphane Pages, Nicolas Chenouard, Ronan Chéreau, Vladimir Kouskoff, Frédéric Gambino, Anthony Holtmaat

## Abstract

The organization of sensory maps in the cerebral cortex depends on experience, which drives homeostatic and long-term synaptic plasticity of cortico-cortical circuits. In the mouse primary somatosensory cortex (S1) afferents from the higher-order, posterior medial thalamic nucleus (POm) gate synaptic plasticity in layer (L) 2/3 pyramidal neurons via disinhibition and the production of dendritic plateau potentials. Here we address whether these thalamocortically mediated responses play a role in whisker map plasticity in S1. We find that trimming all but two whiskers causes a partial fusion of the representations of the two spared whiskers, concomitantly with an increase in the occurrence of POm-driven, N-methyl-D-aspartate receptor (NMDAR)-dependent plateau potentials. Blocking the plateau potentials restores the archetypical organization of the sensory map. Our results reveal a novel mechanism for experience-dependent cortical map plasticity in which higher-order thalamocortically mediated plateau potentials facilitate the fusion of normally segregated cortical representations.

---

Sensory cortices contain functional topographic maps, which can rapidly change in response to training and altered sensory experience (Harding-Forrester and Feldman, 2018). For example, whisker trimming in rodents modifies the proportional representation of spared and trimmed whiskers in the barrel field of the primary sensory cortex (S1) (Feldman, 2009; Feldman and Brecht, 2005; Fox, 2002; Harding-Forrester and Feldman, 2018). This type of cortical map plasticity is thought to be driven by long-term potentiation (LTP) and depression (LTD) of layer (L)4-to-L2/3 and L2/3-L2/3 cortico-cortical (CC) synapses (Clem and Barth, 2006; Clem et al., 2008; Feldman, 2009; Finnerty et al., 1999), as well as by changes in intrinsic neuronal properties and homeostatic mechanisms balancing the loss of surrounding sensory inputs (Gainey and Feldman, 2017; Li et al., 2014). In addition, whisker trimming weakens feed-forward inhibition of L2/3 pyramidal neurons (Gambino and Holtmaat, 2012; House et al., 2011; Jiao et al., 2006), and even may, similarly to monocular deprivation, evoke pruning of inhibitory synapses (Chen et al., 2011; Keck et al., 2011; van Versendaal et al., 2012). Disinhibition could also serve a role in homeostasis by increasing whisker-evoked neuronal spiking (Li et al., 2014), and gate synaptic plasticity (Gambino and Holtmaat, 2012).

Thalamo-cortical (TC) synapses may play a direct or facilitating role in cortical map plasticity. TC axons have been shown to remain plastic throughout life and to be affected by modifications of sensory experience (Jamann et al., 2018; Oberlaender et al., 2012; Wimmer et al., 2010a; Yu et al., 2012). Trimming a subset of whiskers causes a decrease in TC-innervation of deprived but not of spared barrels (Oberlaender et al., 2012; Wimmer et al., 2010a), and sensory learning may induce plasticity of a subset of TC synapses (Audette et al. 2019). However, the relative contributions of CC and TC synaptic plasticity, and how they interact during cortical map plasticity is not known. Moreover, the role of TC synapses may be intricate since different cortical layers receive inputs from diverse thalamic origins, each with distinctive properties.

Sensory information from the whiskers is transmitted to S1 by two main and well-segregated TC projections (Alloway, 2008; Feldmeyer, 2012; Wimmer et al., 2010b). The lemniscal pathway relays sensory information to L5b, L4, and L3 neurons through the ventral posteromedial (VPM) nucleus of the thalamus (Feldmeyer, 2012). The paralemniscal pathway provides a complementary and non-overlapping source of inputs mainly terminating in L5a and L1 that arise from the higher-order posteromedial (POm) nucleus of the thalamus. While the VPM is viewed as the main hub for whisker tactile information to S1, the exact function of the POm in this cortical area remains unclear (Deschênes et al., 2005; Sherman, 2017). Neurons in the POm have broad receptive fields (Diamond et al., 1992; Veinante and Deschênes, 1999), and their axons have extensive arborizations in S1, distributed over multiple barrel-related columns (Feldmeyer, 2012; Jones, 2000; Ohno et al., 2012). POm axons connect to distal pyramidal cell dendrites as well as to various interneurons (Audette et al., 2018; Jouhanneau et al., 2014; Mease et al., 2016; Petreanu et al., 2009; Viaene et al., 2011; Williams and Holtmaat, 2019; Zhang and Bruno, 2019). The large extent of their projections together with their broad receptive fields suggests that POm neurons provide more generalized information to S1 as compared to VPM.

POm projections to S1 mediate whisker-evoked NMDAR-dependent plateau potentials and facilitate whisker-evoked LTP in L2/3 pyramidal neurons (Gambino et al., 2014), which may depend on a combined excitation and disinhibition (Williams and Holtmaat, 2019). In addition, POm projections themselves display plasticity during sensory learning (Audette et al., 2018). Altogether, this suggests that POm projections to S1 could play a distinctive role in the refinement of cortical maps. Here, we investigate the relationship between cortical remapping and POm-mediated plateau potentials upon whisker sensory deprivation. We use a paradigm in which all whiskers were trimmed except from a pair of neighboring ones (dual-whisker experience, DWE). Using intrinsic optical imaging we first confirm electrophysiology studies which showed that DWE causes the representation of the two spared whiskers to partly fuse (Armstrong-James et al., 1994; Diamond et al., 1993a, 1994). We then show that this plasticity is associated with an increase in dendritic plateau potentials, which is dependent on inputs from the POm. The pharmacological removal of the plateau potentials causes the fused whisker representations to segregate, back to an organization seen in naïve mice. Altogether, our results reveal a novel mechanism for rapid experience-dependent cortical map plasticity, which consists of an increased contribution of dendritic plateau potentials that are associated with inputs from higher-order thalamic neuron. This, in turn, may enhance the level of non-specific sensory input and facilitate subsequent synaptic plasticity events that have been shown to underlie cortical map reorganization (Clem et al., 2008; Feldman, 2009; Feldman and Brecht, 2005; Fox, 2002).

## RESULTS

### Dual-whisker experience reshapes whisker-evoked intrinsic optical signals in S1

Single unit and whole cell recordings have shown that DWE causes the functional representation of the spared whiskers in S1 to merge (Diamond et al., 1993b, 1994; Armstrong-James et al., 1994; Feldman, 2009; Feldman and Brecht, 2005; Gambino and Holtmaat, 2012). Intrinsic optical signal (IOS) imaging which is a proxy of whisker-evoked population activity (Grinvald et al., 1986; Cardoso et al., 2012) can potentially quantify such changes at the mesoscale level in a quasi-noninvasive manner (Drew and Feldman, 2009; Polley et al., 1999; Schubert et al., 2013). Here, we used IOS imaging to measure DWE-evoked plasticity of whisker representations in S1.

Mice were separated in two groups. One group was exposed to a brief period of DWE (2-4 days) by clipping all whiskers except C1 and C2, while for the control group all whiskers were left intact to allow full whisker experience (FWE). We used IOS to assess the spatial representation of the C1 and C2 whiskers in S1 under urethane anesthesia (**Figure 1A**). For each mouse, 100 ms-long imaging frames were acquired through the skull before (frames 1-10), during (frames 11-20), and after (frames 21-50) a 1-s long train (8 Hz) of single whisker deflections (**Figure 1A**). The whisker-evoked response area and the corresponding center were then computed by a statistical comparison of the averaged baseline (frames 1-10) and whisker-evoked (frames 19-28) IOS over at least 10 successive trials. This was done by using a pixel-by-pixel paired *t*-test as previously described (Schubert et al., 2013). For each whisker, the resulting *t*-value map was low-pass filtered with a Gaussian kernel (200 μm full width at half maximum) and thresholded (*t*-value = −2). Only pixels with a *t*-value below the threshold were included into the stimulus-evoked response area (**Figure 1B**). The peak of the response area was given by the minimum in the *t*-value distribution. The Euclidian distance between the peaks of the C1 and C2 response areas was used to determine the distance between the two whisker representations (hereafter termed whisker representation distance [WRD]) as a function of time after deprivation (Schubert et al., 2013) (**Figure 1B-D**).

**Figure 1:**
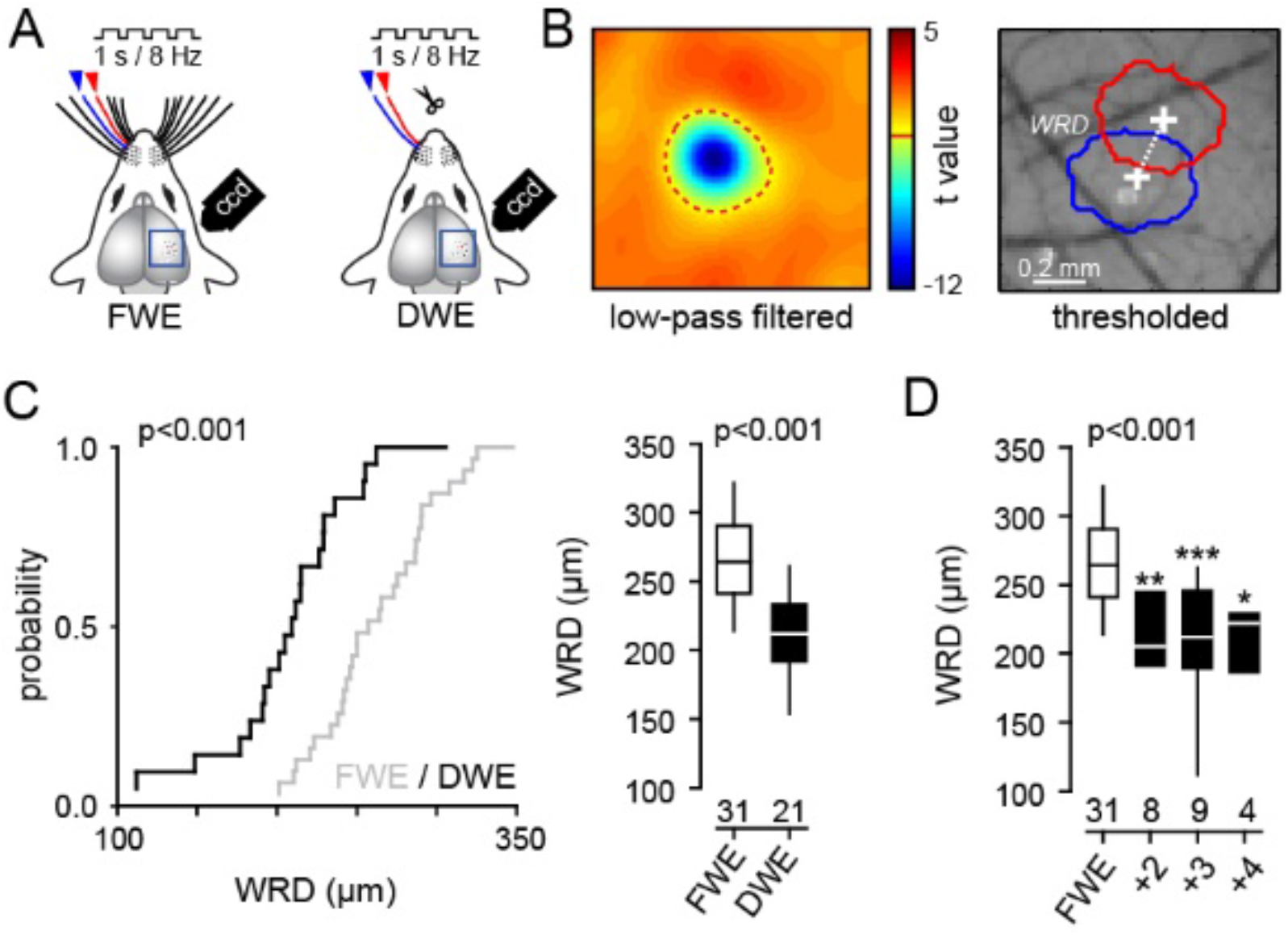
IOS detects DWE-evoked plasticity of whisker representation in S1. **A)** Schematic of whisker trimming and IOS recording. **B)** *Left*, low-pass filtered *t*-value map. Only pixels with a *t*-value lower than −2 are included into the responding area (red dotted line). *Right*, whisker C1 and C2 responding areas. *WRD,* Euclidian distance between the peaks of the C1 and C2 response areas. **C)** *Left*, cumulative distribution of whisker representation distance (WRD) in control mice (FWE) and upon DWE. *Right*, median (± interquartile range) WRD. Number of recorded mice is indicated below. **D)** median (± interquartile range) WRD as a function of deprivation duration (in days).

We found that the WRD was significantly decreased in DWE mice as compared to control mice (FWE: 266 ± 3.7 μm, n=31; DWE: 212 ± 4.1 μm, n=21; *p<0.001*) (**Figure 1C**). This indicates that DWE narrows the distance between the maximally responding populations of neurons, which is in line with the observed merging of whisker representations at the level of neuronal spiking (Armstrong-James et al., 1994; Diamond et al., 1993b, 1994; Drew and Feldman, 2009; Wallace and Sakmann, 2008). We found that increasing the duration of DWE had no further effect on the WRD (**Figure 1D**), indicating that the merging had reached a maximum within 2 days and remained stable for at least 4 days. Importantly, it occurred at a time at which no alterations in activity of layer 4 granular neurons have been observed (Diamond et al., 1993b, 1994), suggesting that the changes in IOS primarily originate in alterations of neural activity within L2/3 (Diamond et al., 1994; Glazewski and Fox, 1996; Stern et al., 2001).

### DWE increases NMDAR-mediated dendritic plateau probabilities and long-latency action potentials in L2/3 pyramidal neurons

Next, we performed whole-cell recordings of L2/3 pyramidal neurons *in vivo* in the C2 barrel-related column while deflecting either the principal (PW, C2) or surrounding (SW, C1) whisker, in FWE mice or after DWE (**Figure 2**). In accordance with previous reports (Armstrong-James et al., 1993; Gambino and Holtmaat, 2012; Petersen et al., 2003; Wilent and Contreras, 2004), single principal whisker deflections typically evoked compound postsynaptic potentials (PSPs) that contained short and long-latency components (**Figure 2A, B**). The latter might represent dendritic NMDAR-mediated potentials that spread towards the soma (Gambino et al., 2014; Palmer et al., 2014). Short-latency PSPs were reliably evoked in successive trials, with a peak amplitude that was always higher upon PW deflections as compared to SW deflections (PW: 9.95 ± 0.9 mV, n=33; SW: 6.67 ± 0.7 mV, n=31; *p<0.001*) (**Figure S1**). In contrast, long-latency PSPs occurred with variable probabilities (**Figure 2B**). We extracted these long-latency PSPs as previously described (Gambino et al., 2014). Briefly, for each whisker deflection, the relationship between the PSP half-peak amplitude and the average membrane potential between 50 and 100 ms after the onset reveals two distinct clusters of sensory-evoked PSP (**Figure 2B, C**). Cluster 1 was defined by an index < 0, which consisted of short latency PSPs that quickly returned to the resting membrane potential. Cluster 2 was defined by an index > 0 (**Figure 2C**), which consisted of compound PSPs containing both short and long-latency components. The long-latency component of the PSPs in cluster 2 was obtained by subtracting the peak-scaled PSP average of cluster 1 from the PSP average of cluster 2 (**Figure 2D**). It was previously shown that these late components disappears when NMDAR conductances are blocked, and thus represent dendritic plateau potentials (Gambino et al., 2014).

**Figure 2:**
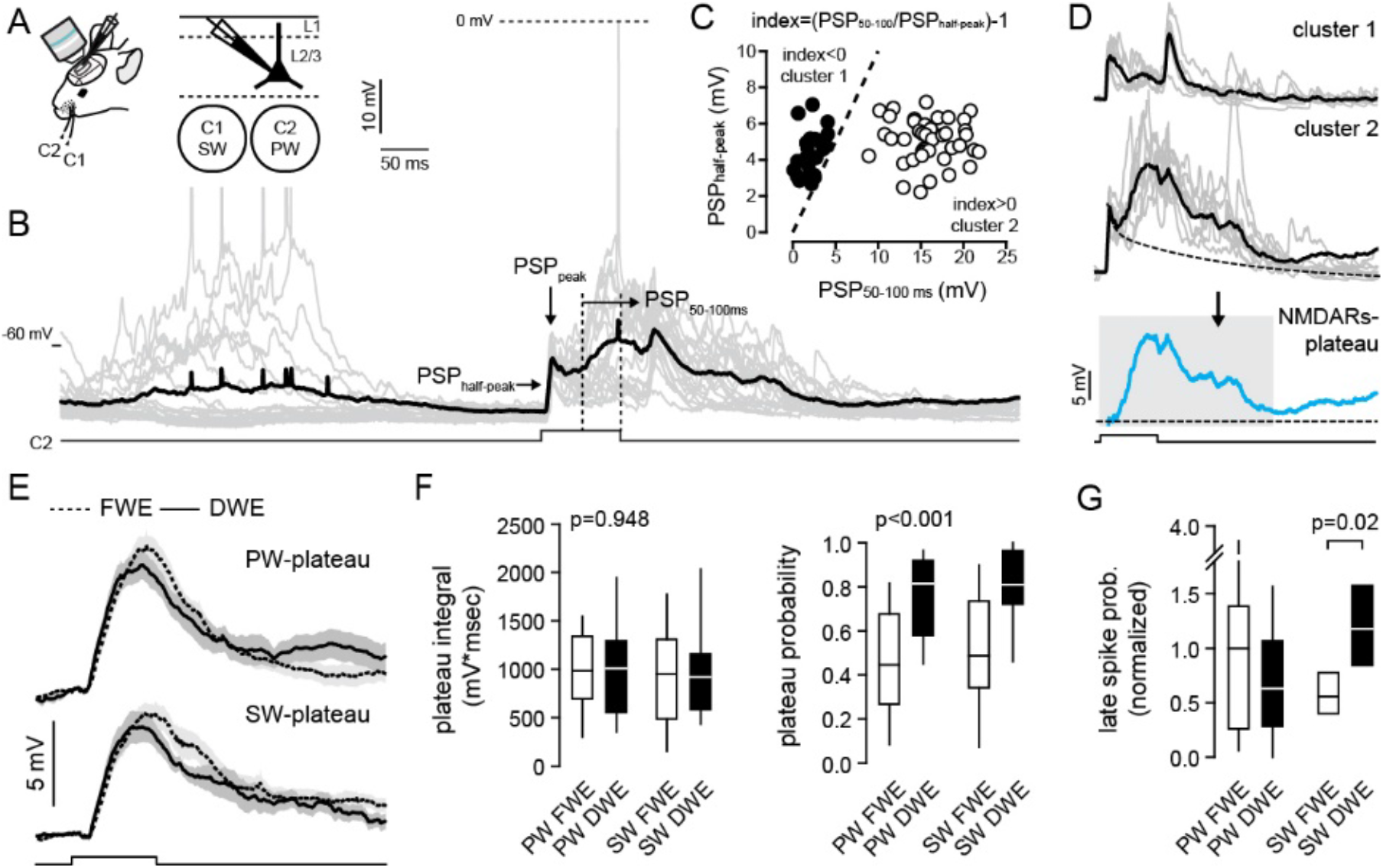
DWE increases plateau potential probabilities. **A)** Schematic of recordings in L2/3 cells *in vivo* in the C2 barrel-related column. **B)** Single-cell examples of principal (C2) whisker-evoked responses (grey, single trials traces; dark, averaged trace). Square pulse lines, C2 whisker deflection (100 ms). **C)** For each trial, the relationship between the PSP half-peak amplitude and the average membrane potential between 50 and 100 ms after the onset reveals two distinct clusters. Dotted line represents the identity line. **D)** Cluster 1 is defined by an index < 0 and consists of PSP containing only a short latency PSP that quickly returns to the resting membrane potential. Cluster 2 is defined by an index > 0 and consists of compound PSP with short and long-latency components. The long-latency component of the PSP depends on NMDAR (Gambino et al., 2014). For each cell, the NMDAR-plateau potential (bottom) is derived by subtracting mean cluster 1 response from mean cluster 2 response. The integral of the plateau potential is measured from 0 to 300 ms (grey box). **E)** Plateau potential grand average (all recorded cells averaged) ± sem, evoked by the PW (top) and SW (bottom), in control mice (dotted line, FWE) and upon DWE (solide line). Square pulse lines, whisker deflection (100 ms). **F)** Median (± interquartile range) plateau potential integral (left) and probability (right). **G)** Mean (± interquartile range) late spike probability (normalized to the spiking probability measured in control mice upon PW stimulation). For E-G, PW and SW correspond to C2 and C1 whiskers respectively.

We made comparisons between the extracted NMDAR-mediated plateaus elicited by the PW and the SW in FWE and DWE mice (**Figure 2E**). In FWE mice, plateau potentials were, in contrast to short-latency PSPs, elicited by both whiskers to a similar extent (PW: 985 ± 87 mV*msec, n=33; SW: 951 ±108 mV*msec, n=31; *p=0.396*) and with similar probabilities (PW: 0.47 ± 0.04, n=33; SW: 0.51 ±0.05, n=31; *p=0.305*). This suggests that NMDAR-mediated plateau potentials are not whisker-specific (Gambino et al. 2014). DWE did not affect the plateau-potential integrals (*p=0.948*) (**Figure 2F**), nor the short-latency PSPs (**Figure S1**). However, DWE did increase the SW/PW ratio of short-latency peak amplitudes, confirming that this paradigm does cause the relative strengthening of SW-associated inputs (Armstrong-James et al., 1994; Diamond et al., 1993a, 1994; Gambino and Holtmaat, 2012) (**Figure S1**). In addition, DWE significantly increased the probability of both the PW and SW-evoked plateau potentials as compared to controls (FWE, PW: 0.47 ± 0.04, n=33; SW: 0.51 ±0.05, n=31; DWE, PW: 0.762 ± 0.04, n=20; SW: 0.786 ±0.05; n=20; *p<0.001*) (**Figure 2F**). In addition, we observed that the NMDAR-mediated plateau potentials occasionally elicited action potentials. These were triggered with long delays after the whisker stimulus (**Figure 2B**), which is consistent with the earlier finding that long-latency spikes in the barrel cortex may depend on NMDARs (Armstrong-James et al., 1993; Salt, 1986). In line with the increased probability of evoked plateau potentials, DWE also increased the probability of SW-evoked long-latency spikes (**Figure 2G**). Collectively, these data indicate that DWE concomitantly increases the probability of whisker-evoked NMDAR-mediated plateau potentials and spikes in L2/3 pyramidal neurons, and merges the cortical representation of the two spared whiskers.

### Increased NMDAR-mediated plateau potential probabilities depend on paralemniscal synaptic input

In naive mice, NMDAR-mediated plateau potentials in L2/3 pyramidal neurons depend on paralemniscal synaptic inputs from the posteromedial nucleus (POm) of the thalamus (Gambino et al., 2014). Here, we hypothesized that the increase in plateau potentials upon DWE is both NMDAR and POm-dependent. First, we confirmed that the plateau potential probability decreased in the presence of the NMDAR open-channel blocker MK-801 (1 mM) inside the intracellular solution (iMK801; DWE/control: 0.762 ± 0.04, n=20; DWE/+iMK801: 0.2 ± 0.07, n=7; *p<0.001*) (**Figure 3B-D**). A local injection of the GABA-A receptor (GABA-AR) selective agonist muscimol into the POm also reduced the probability of plateau potentials (**Figure 3B-D**). This did not occur when the injection was incorrectly targeted within the thalamus (DWE/muscimol in the POm: 0.14 ± 0.04, n=6; DWE/muscimol excluded from the POm: 0.7 ± 0.04, n=6; *p<0.001*) (**Figure 3B-D**). Since these pharmacological interventions may not only affect probabilities but also the magnitude of the plateau potentials, we measured the net plateau strength under all conditions, which is the product of the integral of the potential and its probability for any whisker stimulus-evoked response. This significantly decreased when NMDARs or the activity of the POm were blocked (**Figure 3E, F; Figure S1**), though the integrals of those plateau potentials that remained were unaffected (**Figure S1**). Taken together, our data suggest that DWE facilitates the occurrence of whisker-evoked NMDAR-plateau potentials in L2/3 pyramidal neurons, gated by input from the POm.

**Figure 3:**
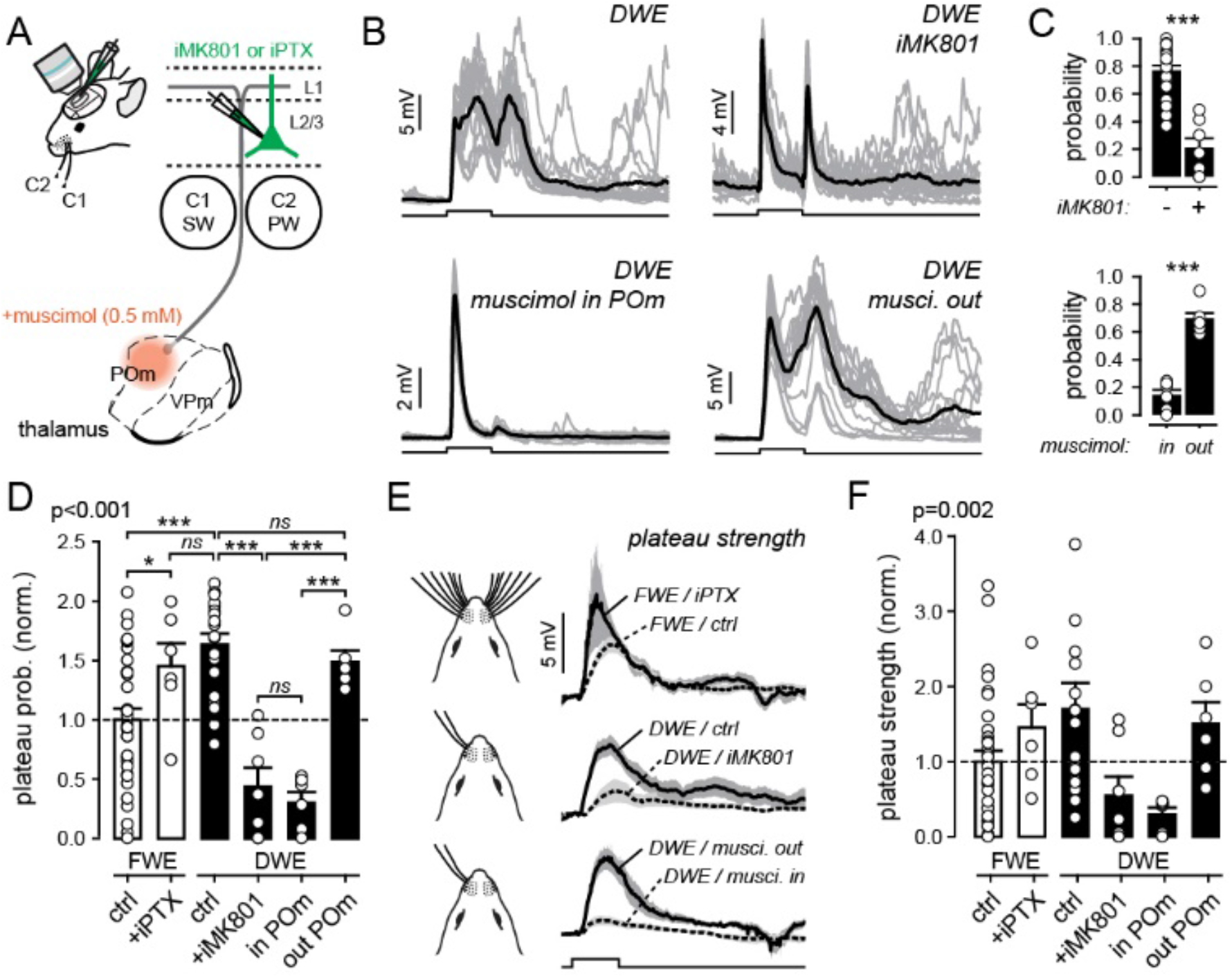
DWE increases the probability of POm-mediated, NMDAR-dependent plateau potentials. **A)** Schematic of the thalamo-cortical circuit and pharmacological experiments. Fluorescent muscimol (0.5 mM) is injected locally in the POm (in POm) or in structures not directly involved in somatosensation (out POm) for controls. The GABA-A receptor antagonist picrotoxin (iPTX, 1 mM) or the NMDAR open-channel blocker MK-801 (iMK801, 1 mM) are applied directly to the intracellular recording solution **B)** Single-cell example of whisker-evoked responses in different conditions. Gray lines, individual trials; Black lines, averaged traces. Square pulse lines, C2 whisker deflection (100 ms) Mean (± sem) plateau potential probability after DWE. Circles, individual cells. **D)** Mean (± sem) plateau potentials in FWE and DWE mice under different pharmacological conditions. Square pulse lines, C2 whisker deflection (100 ms) **E)** Grand average of PW-evoked plateau potential strength (extracted and averaged from all recorded cells) ± sem in control (FWE) and after DWE, under different pharmacological conditions. **F)** Mean (± sem) plateau potentials strength in control (FWE) and after DWE, under different pharmacological conditions, normalized to the mean measured in FWE mice (dashed line).

GABA-AR-mediated inhibition in the barrel cortex shunts excitatory conductances in pyramidal distal dendrites and spines (Koch, 1999; Larkum et al., 1999, 2007; Palmer et al., 2012) and impairs NMDAR-dependent synaptic plasticity (Gambino and Holtmaat, 2012; Williams and Holtmaat, 2019). Thus, inhibitory inputs may gate whisker-evoked NMDAR-mediated plateau potentials (Palmer et al., 2012). To test this we added the GABA-AR antagonist picrotoxin to the intracellular recording solution (iPTX, 1 mM), which has been shown to efficiently suppress whisker-evoked inhibition in pyramidal neurons, probably through the small and local diffusion of the drug in and around the recorded neuron (Gambino and Holtmaat, 2012; Yazaki-Sugiyama et al., 2009). The GABA-AR block increased the probability of plateau potentials in FWE mice (FWE/ctrl: 0.46 ± 0.04, n=33; FWE/+iPTX: 0.67 ± 0.09, n=6; *p=0.03*) (**Figure 3D; Figure S2**). Importantly, the probability increased to a level that was similar to DWE mice (DWE/ctrl: 0.762 ± 0.04, n=20; FWE/+iPTX: 0.67 ± 0.09, n=6; *p=0.4*) (**Figure 3D**). Similar results were obtained when plateau potential strength was considered (**Figure 3E, F**), suggesting that GABA-AR inhibition is involved in the gating of POm-dependent NMDAR-mediated plateau potentials. Altogether, our results indicate that the DWE-evoked increase in plateau potential probabilities depends on NMDAR and paralemniscal inputs, possibly facilitated by disinhibition.

### Relationship between whisker-evoked plateau potentials and functional map reorganization

In FWE mice the PW and SW drove plateau potentials with similar probabilities (**Figure 2F**) (Gambino et al. 2014), suggesting that they are whisker non-specific. The plateau potential occurrence probabilities were enhanced upon DWE (**Figure 2F**), which increased the spike rates for SW deflections (**Figure 2G**). Together, this implies that the decrease in WRD as seen in IOS imaging (**Figure 1**) might depend on plateau potential-mediated mechanisms. To further explore the relationship between plateau potentials and the merging of whisker representations, we first measured the average plateau potential strength for PW and SW over DWE and FWE mice. This shows that DWE significantly increased the PW but not the SW-evoked plateau strength (PW, FWE: 0.51 ± 0.07, n=33; PW, DWE: 0.87 ± 0.17, n=20; *p=0.047;* SW, FWE: 0.56 ± 0.09, n=31; SW, DWE: 0.81 ± 0.13, n=20; *p=0.088*) (**Figure 4A-D**). Then we expressed the strength of PW- and SW-evoked plateau potentials as a function of the distance between the spared whisker-evoked IOS centers in DWE mice (**Figure 4E**). We observed that for each of the two spared whiskers, the level of plateau strength negatively correlated with the WRD (PW, r^2^=0.47, *p<0.01*; SW, r^2^=0.75, *p<0.001*). Compared to FWE mice, both PW and SW deflections induced stronger plateau potentials in DWE mice with the smallest WRD (WRD < λ; PW FWE: 0.51 ± 0.07, n=33; PW DWE: 1.3 ± 0.3, n=9; *p=0.007*; SW FWE: 0.56 ± 0.09, n=31; SW DWE: 1.07 ± 0.25, n=9; *p=0.033*). This indicates that the merging of the spared whisker representations is tightly coupled to the plateau strength (**Figure 4E**).

**Figure 4:**
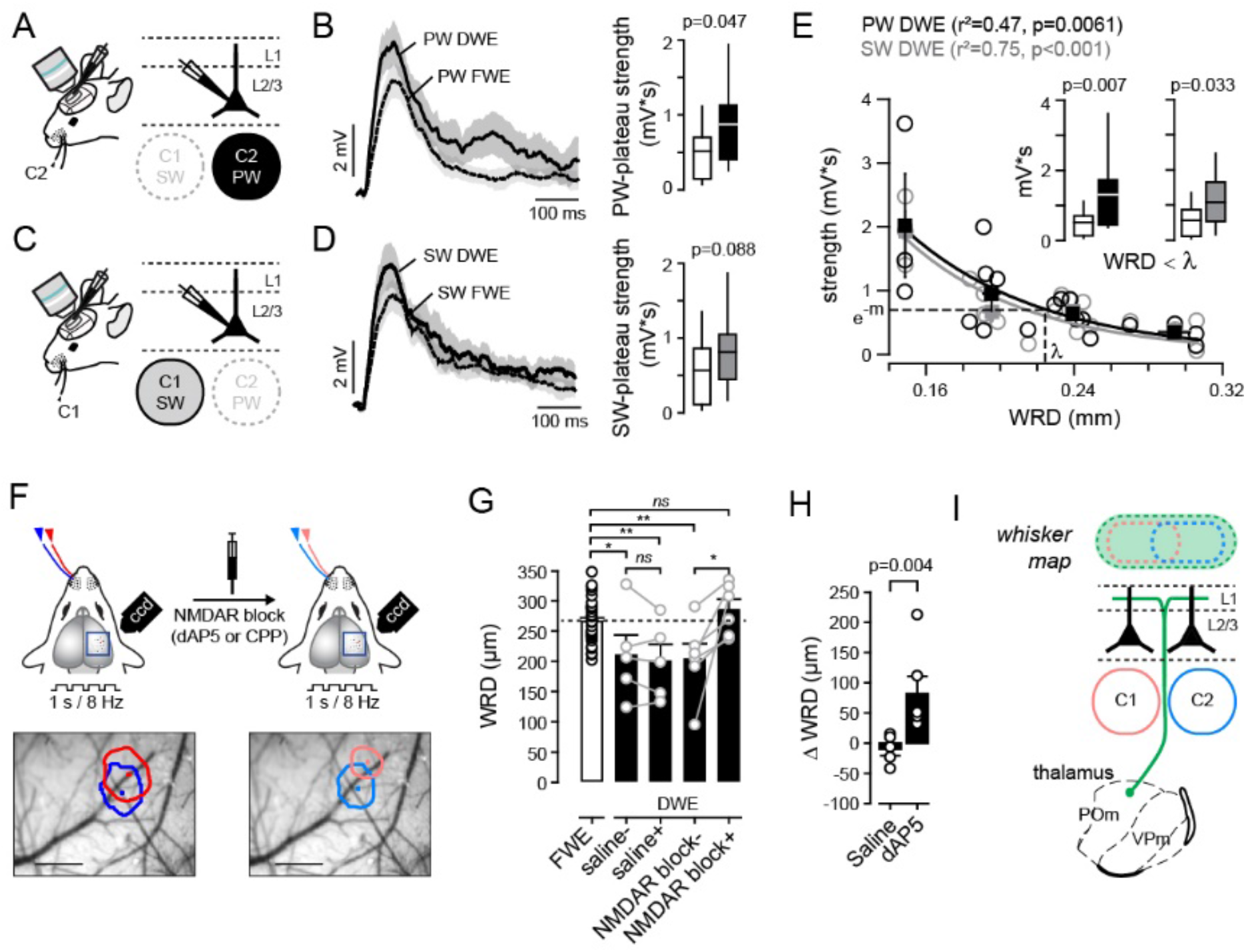
Blocking NMDAR-dependent plateau potentials restores the IOS map. **A)** Schematic of recordings in L2/3 cells *in vivo*. PSP are evoked by the principal whisker (C2). **B)** *Left,* grand average of PW-evoked plateau potential strength (extracted and averaged from all recorded cells) ± sem in control (FWE) and after DWE. *Right*, Median (± interquartile range) plateau potential strength. **C-D)** Same presentation as in (**A-B**) but for SW-evoked plateau strength. **E)** Relation between WRD and PW-(black) and SW-(gray) plateau strength. Circles, individual cells; Squares, averages. *Inset*, median (± interquartile range) plateau potential strength, for WRD < λ, in control naïve mice (white) and after DWE (black, PW; gray, SW). **F)** Schematic of experimental protocol. IOS is obtained in DWE mice, before (NMDAR block−) and after (NMDAR block+) the suppression of NMDAR conductances by applying dAP5 or CPP (or saline for controls). **G)** Mean (± sem) WRD in control (FWE) and after DWE, under different pharmacological conditions. **H)** ΔWRD in DWE mice following saline or dAP5 application. Blocking NMDAR conductance significantly increases the inter-barrel distance, and thus restore the intrinsic whisker map. **I)** NMDAR-dependent, POm-driven plateau potentials participates in the functional fusion of spared whiskers cortical representations upon DWE.

We then measured the WRD for the two spared whiskers upon DWE before and after the topical application of the NMDAR antagonist dAP5 (or saline for control) (**Figure 4F**). First, we confirmed that DWE decreased the WRD (FWE: 265.7 ± 36 μm, n=31; DWE/saline-: 210 ± 39 μm, n=5; DWE/dAP5-: 203 ± 25 μm, n=6) (ctrl *vs.* DWE/saline-, *p=0.019*; ctrl *vs.* DWE/dAP5-, *p=0.005*) (**Figure 4G**). Whereas the WRD remained unchanged upon an injection of saline, it significantly increased upon the pharmacological suppression of the NMDAR conductance (DWE, NMDAR block-: 203 ± 25 μm, n=6; NMDAR block+: 285 ± 17 μm, n=6; *p=0.031;* ΔWRD: +82 ± 29 μm; DWE, saline-: 210 ± 39 μm; saline+: 200 ± 28 μm, n=5; *p=0.398;* ΔWRD: −9 ± 10 μm) (**Figure 4G, H**). Importantly, upon blocking NMDARs the distances between the whisker-evoked IOS areas increased to levels that were observed in FWE mice (FWE: 265.7 ± 36 μm, n=31; DWE/ NMDAR block+: 285 ± 17 μm, n=6) (**Figure 4G**). Altogether, our results suggest that the increase in plateau potential strength participates in the DWE-evoked fusion of spared whisker representations in S1 (**Figure 4I**).

## DISCUSSION

We used a dual whisker experience (DWE) paradigm in mice to investigate mechanisms of cortical map plasticity in S1. In this paradigm, the trimming of all but two adjacent whiskers causes the spared whiskers to increase their excitatory drive of neurons in the neighboring spared barrel column but not in the deprived areas (Armstrong-James et al., 1994; Diamond et al., 1993a, 1994). This plasticity is more modest as compared to a single whisker experience paradigm, in which the expansion of the spared whisker representation extends far into deprived cortical areas (Feldman, 2009; Glazewski et al., 1996). Imaging of intrinsic optical signals, which has been described to correlate with sensory-evoked spiking on a ~100 μm spatial scale (Ts’o et al., 1990), readily captures single-whisker map plasticity (Polley et al., 1999). Using IOS imaging, we found that the distance between the centers of the spared whisker’s cortical representations narrows significantly (**Figure 1**). This recapitulates the results of extracellular recordings in that it detects the mutual expansion of spared surround whisker-evoked neuronal population activity in both spared barrel columns (Feldman, 2009; Feldman and Brecht, 2005). Thus, our data confirm that IOS imaging has sufficient resolution to visualize even subtle forms of map plasticity such as found under anesthesia upon DWE (Li et al., 2014).

Whisker map changes upon DWE are thought to be driven primarily by modulated activity of L2/3 and governed by Hebbian forms of plasticity (Armstrong-James et al., 1994; Diamond et al., 1993a, 1994). Remarkably, only a small fraction of L2/3 pyramidal neurons in S1 discharges action potentials in response to a single whisker deflection (Wolfe et al., 2010; Kock et al., 2009). This implies that the map changes as observed using IOS imaging are constituted by alterations in a sparsely spiking population of neurons. We found that DWE does not only change the ratios of PW and SW-driven short latency spikes, but also promotes the generation of SW-driven long-latency spikes (**Figure 2**) (Armstrong-James et al., 1993). Since IOS integrate activity over relatively long timespans, these long-latency spikes could have significantly contributed to the reduced distance between the spared whisker representations. This implies that IOS map changes depend on the increased occurrence of plateau potentials, since they primarily drove the long-latency spikes. Indeed, a block of NMDARs after DWE decreased plateau potential probabilities and restored the distance between neighboring spared whisker-evoked IOS (**Figures 3 and 4**). Altogether, this strongly suggests that the DWE-evoked cortical map changes as observed using IOS are associated with an increased occurrence of dendritic plateau potentials. It is tempting to speculate that the IOS map changes not only depend on somatic short and long-latency spikes, but are also directly generated by the subthreshold plateau potentials, which are driven by local active mechanisms in apical dendrites and are accompanied by substantial ion flux (Antic et al., 2010; Gambino et al., 2014; Larkum et al., 2009; Major et al., 2008; Palmer et al., 2014). Since IOS may strongly depend on ion-related water movements (Vincis et al., 2015), such strong ion fluxes could have contributed to the DWE-evoked map changes in our experiments.

Multiple mechanisms could explain the increase in whisker-evoked plateau potentials. Here (**Figure 3**), as in previous work (Gambino et al., 2014), we show that plateau potentials in the barrel cortex depend in part on activity of the POm division of the thalamus. POm neurons send dense axonal projections to L1 in S1 (Ohno et al., 2012; Wimmer et al., 2010b), where they spread out over multiple barrel columns and make synaptic contacts with apical dendrites of numerous L2/3 pyramidal neurons (Bureau et al., 2006; Feldmeyer, 2012; Gambino et al., 2014; Jones, 2000; Jouhanneau et al., 2014; Ohno et al., 2012; Petreanu et al., 2009; Sermet et al., 2019; Zhang and Bruno, 2019). Interestingly, neurons located in this higher-order thalamic nucleus have large receptive fields (Diamond et al., 1992; Jouhanneau et al., 2014; Masri et al., 2008), and the POm input-recipient L2/3 neurons are characterized by large and short-latency responses to multiple whisker deflections (Jouhanneau et al., 2014). Thus, the increase in plateau potentials upon DWE could point to an increased activity of POm neurons projecting to L1. POm neurons receive powerful inhibitory inputs from the zona incerta (ZI) (Lavallée et al., 2005; Trageser and Keller, 2004), which might in turn affect the function and activity of POm depending on the strength of this inhibition, notably during pathological conditions (Masri et al., 2009). In addition, the ZI-POm connections are strongly modulated by the release of neuromodulators such as acetylcholine (Ach), raising the possibility that POm activity could be strongly gated by arousal (Masri et al., 2006; Trageser et al., 2006).

Another mechanism for the increase in plateau potentials may include the additional cholinergic effects on dendritic computational properties. Acetylcholine promotes the generation of long-lasting dendritic plateau potentials (Williams and Fletcher, 2019) that could eventually facilitate the plasticity of TC projections (Dringenberg et al., 2007). On the other hand, accumulating evidence suggests that plateau potentials are strongly and specifically controlled by dendrite-targeted inhibition (Larkum, 2013; Palmer et al., 2012). Here, we found that locally blocking GABA-AR-mediated inhibition dramatically increased the occurrence of whisker-evoked plateau potentials in naive mice. Mechanistically, this modulation of dendritic excitability could be driven by the inhibition of interneurons that specifically shunt synaptic inputs from TC projections (Koch, 1999; Kubota et al., 2007), and/or by the stimulation of TC-mediated disinhibitory motifs (Audette et al., 2018; Williams and Holtmaat, 2019). In line with these possibilities, it is becoming increasingly clear that sensory map plasticity depends on intricate changes in inhibitory and disinhibitory circuits (Gainey and Feldman, 2017; Gambino and Holtmaat, 2012; Harding-Forrester and Feldman, 2018; Li et al., 2014).

What could be the consequences of the increase in dendritic plateau potentials upon alterations of sensory experience? Previous works show that plateau potentials are strong drivers of synaptic LTP (Brandalise et al., 2016; Gambino et al., 2014; Golding et al., 2002), which is intimately associated with map plasticity in the barrel cortex (Feldman, 2009; Feldman and Brecht, 2005; Glazewski et al., 2000). Moreover, TC inputs from POm can also drive cortical LTP through disinhibition (Williams and Holtmaat, 2019). Thus, upon DWE, an increase of POm-originating inputs or activity thereof might facilitate LTP in L2/3 pyramidal neurons by evoking disinhibition and plateau potentials, both of which generate favorable conditions for the integration, stabilization and strengthening of relevant synaptic inputs (Holtmaat and Caroni, 2016). Interestingly, only pyramidal neurons located in the supragranular (L2/3) and infragranular (L5) layers of the spared barrels, but not in L4, rapidly increase their activity in response to changes in sensory experience (Diamond et al., 1993b, 1994). This observation has led to the hypothesis that, in adult animals, plasticity occurs first in L2/3 and L5 (Feldman and Brecht, 2005). Thus, it is conceivable that L2/3 and L5 pyramidal neurons rapidly respond to DWE by a combination of whisker-nonspecific POm input and disinhibition, resulting in increased plateau potentials which subsequently may lead to elevated levels of synaptic plasticity.

Functional studies *in vivo* have highlighted the pivotal role of dendritic non-linear events in sensory-evoked spiking and plasticity (Cichon and Gan, 2015; Du et al., 2017; Gambino et al., 2014; Palmer et al., 2014), as well as the control of active behavior and perceptual discrimination (Takahashi et al., 2016; Xu et al., 2012). It will be interesting in future studies to dissect the relationship between higher-order thalamic inputs to cortex and experience-dependent synaptic and map plasticity.

## ACKNOWLEDGEMENTS

We thank all the members of the Gambino and Holtmaat laboratories for technical assistance and helpful discussions. This project has received funding from (to FG): the European Research Council (ERC) under the European Union’s Horizon 2020 research and innovation program (grant agreement n° 677878) and the FP7 Marie-Curie Career Integration program (grant agreement n° 631044); the ANR JCJC (grant agreement n° 14-CE13-0012-01), the University of Bordeaux (Initiative of Excellence senior chair 2014); (to AH): the Swiss National Science Foundation (grants 31003A-153448, 31003A_173125, CRSII3_154453, and NCCR Synapsy 51NF40-158776), and a gift from a private foundation with public interest through the International Foundation for Research in Paraplegia (chair Alain Rossier).

## AUTHOR CONTRIBUTION

SP, NC, RC, VK, and FG performed the experiments and analyzed the data. FG and AH conceived the studies, supervised the research and wrote the manuscript with the help from co-authors.

## DECLARATION OF INTERESTS

The authors declare no competing financial interests.

---

**Figure S1.**
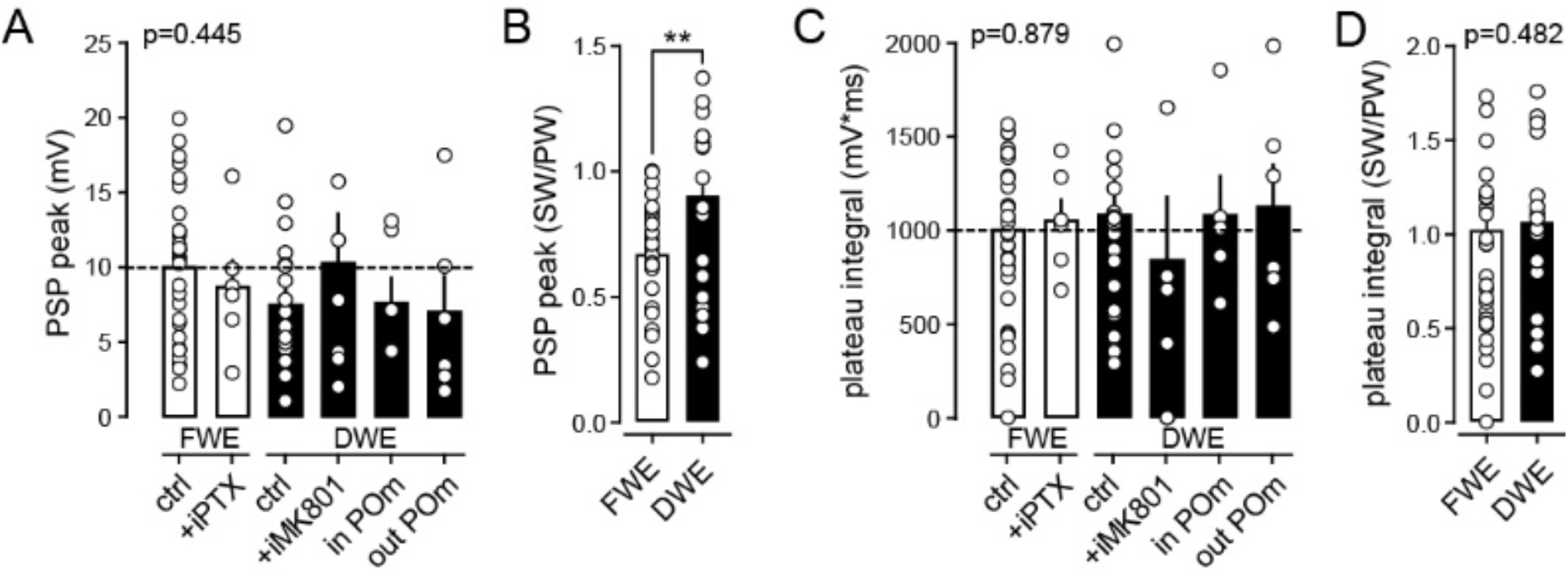
Comparisons of PW-evoked PSP peak and plateau potentials between all conditions. **A)** Mean (± sem) PSP peak amplitude in control mice (FWE) and after DWE, under different pharmacological conditions. **B)** SW/PW ratio of PSP peak amplitudes. **C)** Mean (± sem) PSP plateau potential integral in control mice (FWE) and after DWE, under different pharmacological conditions. **D)** SW/PW ratio of plateau potentials integrals.

**Figure S2.**
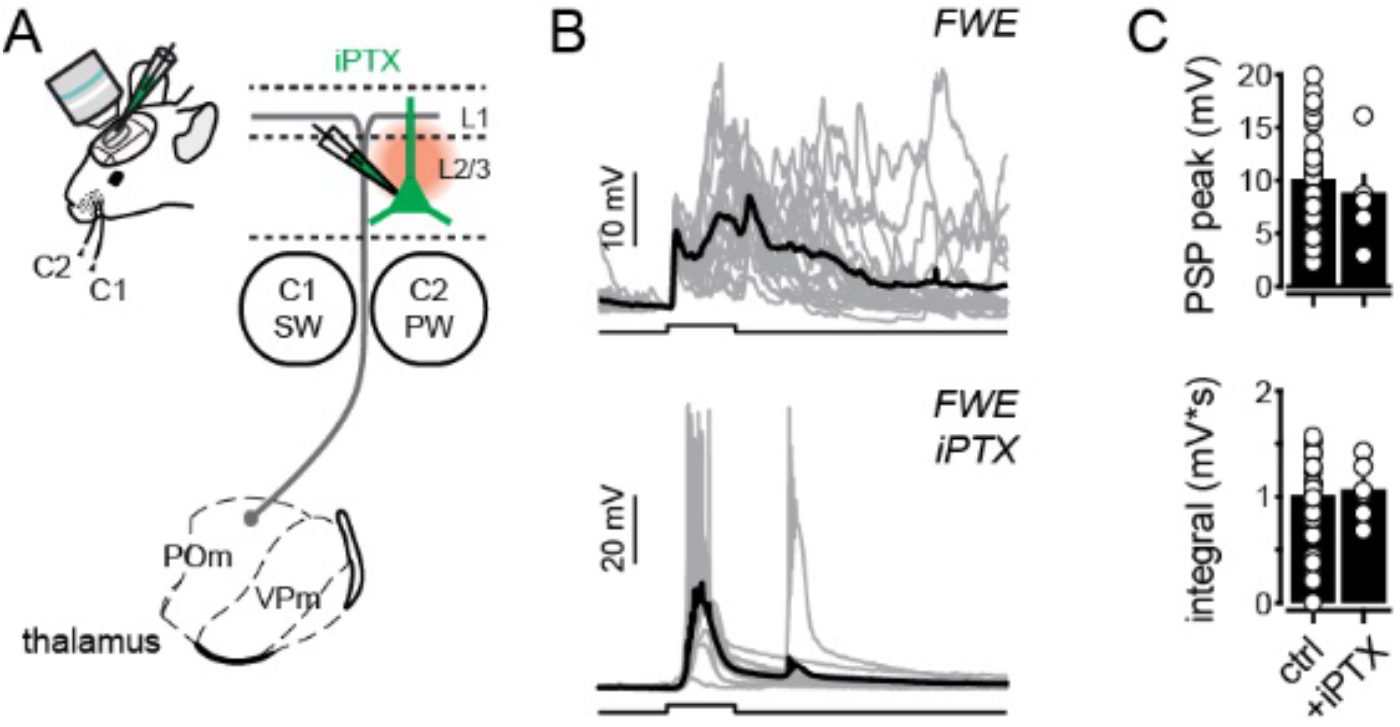
Effect of GABA-ARs blockade in L2/3 pyramidal neuron *in vivo*. **A)** Schematic of the thalamo-cortical circuit and pharmacological experiments. The GABA-A receptor antagonist picrotoxin (iPTX, 1 mM) is applied directly to the intracellular recording solution **B)** Single-cell example of whisker-evoked responses in controls (top, FWE) and during GABA-AR blockage (bottom, FWE, iPTX). Gray lines, individual trials; Black lines, averaged traces. Square pulse lines, C2 whisker deflection (100 ms) **C)** Mean (± sem) PW-evoked PSP peak amplitude (top) and integrals plateau potential probability. Circles, individual cells.

## METHODS

All experiments were performed in accordance with the Guide for the Care and Use of Laboratory Animals (National Research Council Committee (2011): Guide for the Care and Use of Laboratory Animals, 8th ed. Washington, DC: The National Academic Press.) and the European Communities Council Directive of September 22th 2010 (2010/63/EU, 74), as well as the Federal Food Safety and Veterinary Office of Switzerland and in agreement with the veterinary office of the Canton of Geneva (licence numbers GE/28/14, GE/61/17, and GE/74/18). Experimental protocols were approved by the institutional ethical committee guidelines for animal research (N°50DIR_15-A) and by the French Ministry of Research (agreement N°18892). We used male C57BL6/J 5- and 6-weeks old mice from Charles River that were housed with littermates (3 mice per cage) in a 12-h light-dark cycle. Cages were enriched with tunnels. Food and water were provided *ad libitum*.

### Intrinsic optical imaging

Intrinsic optical signals were obtained through the intact skull using a light guide system with a 700 nm (bandwidth of 20 nm) interference filter and stable 100-W halogen light source, as previously described (Schubert et al., 2013). Briefly, Isoflurane (4% with ~0.5 l/min O_2_) combined with an i.p. injection of urethane (1.5 g/kg, in lactated ringer solution containing in [mM] 102 NaCl, 28 Na L Lactate, 4 KCl, 1.5 CaCl_2_) was used to induce anesthesia and prolonged by supplementary urethane (0.15 g.kg^−1^) if necessary. To prevent risks of inflammation, brain swelling and salivary excretions, 40 μl of dexamethasone (Dexadreson, 0.1 mg/ml, i.m.) and glycopyrrolate (Robinul-V, 0.01 mg/kg, s.c.) were injected before the surgery. Adequate anesthesia (absence of toe pinch and corneal reflexes, and vibrissae movements) was constantly checked and body temperature was maintained at 37°C using a heating-pad positioned underneath the animal. Ophthalmic gel was applied to prevent eye dehydration. Analgesia was provided as described for viral injection (with lidocaine and buprenorphine). After disinfection of the skin (with modified ethanol 70% and betadine), the skull was exposed and a ~3mm plastic chamber was attached to it above the prefrontal cortex using a combination of super glue (Loctite) and dental acrylic and dental cement (Jet Repair Acrylic, Lang Dental Manufacturing).

The head of the animal was stabilized using a small stereotaxic frame and the body temperature kept constant with a heating pad. An image of the surface vascular pattern was taken using a green light (546 nm interference filter) at the end of each imaging session. Images were acquired using the Imager 3001F (Optical Imaging, Mountainside, NJ) equipped with a large spatial 602 × 804 array, fast readout, and low read noise charge-coupled device (CCD) camera. The size of the imaged area was adjusted by using a combination of two lenses with different focal distances (upper lens: Nikon 135 mm, f2.0; bottom lens: Nikon 50 mm, f1.2). The CCD camera was focused on a plane 300 μm below the skull surface. Images were recorded at 10 Hz for 5 sec, with a spatial resolution of 4.65 μm/pixel comprising a total area of 2.9 × 3.7 mm^2^. Whisker C2 was deflected back and forth (20 stimulations at 8 Hz for 1 sec.) using a glass-capillary attached to a piezoelectric actuator (PL-140.11 bender controlled by an E-650 driver; Physik Instrumente) triggered by a pulse stimulator (Master-8, A.M.P.I.). Each trial consisted of a 1 sec. of baseline period (frames 1-10), followed by a response period (frames 11-20) and a post-stimulus period (frames 21-50). Inter-trial intervals lasted 20 sec to avoid contamination of the current intrinsic optical signal by prior stimulations. Intrinsic signals were computed by subtracting each individual frame of the response period by the average baseline signal. The obtained intrinsic signal was overlapped with the vasculature image using ImageJ software (Schneider et al., 2012) to precisely identify the C2 whisker cortical representation.

### In vivo electrophysiology

#### Whole-cell recordings

After intrinsic optical imaging, a small ~1 × 1 mm craniotomy (centered above the C2 whisker maximum intrinsic optical response) was made using a pneumatic dental drill, leaving the dura intact. Whole-cell patch-clamp recordings of L2/3 pyramidal neurons were obtained as previously described (Gambino et al., 2014). Briefly, high-positive pressure (200–300 mbar) was applied to the pipette (5–8 MΩ) to prevent tip occlusion, when passing the pia. Immediately after, the positive pressure was reduced to prevent cortical damage. The pipette resistance was monitored in the conventional voltage clamp configuration during the descendent pathway through the cortex (until −200 μm from the surface) of 1 μm steps. When the pipette resistance abruptly increased, the 3–5 GΩ seal was obtained by decreasing the positive pressure. After break-in, Vm was measured, and dialysis could occur for at least 5 min before launching the recording protocols. Current-clamp recordings were made using a potassium-based internal solution (in mM: 135 potassium gluconate, 4 KCl, 10 HEPES, 10 Na2-phosphocreatine, 4 Mg-ATP, 0.3 Na-GTP, and 25 μM, pH adjusted to 7.25 with KOH, 285 mOsM), and acquired using a Multiclamp 700B Amplifier (Molecular Devices). Spiking pattern of patched cells was analyzed to identify pyramidal neurons. Offline analysis was performed using custom routines written in Sigmaplot (Systat), IGOR Pro (WaveMetrics) and Matlab (Mathworks).

#### Whisker evoked postsynaptic potentials (PSPs) in down state

Whisker-evoked PSPs were evoked by forth and back deflection of the whisker (100 ms, 0.133 Hz) using piezoelectric ceramic elements attached to a glass pipette ~4 mm away from the skin. The voltage applied to the ceramic was set to evoke a whisker displacement of ~0.6 mm with a ramp of 7-8 ms. The C1 and C2 whiskers were independently deflected by different piezoelectric elements. The amplitudes of the evoked PSPs were more pronounced during down states as opposed to the up states. Therefore, to facilitate comparisons of PSPs under different conditions, analysis was confined to peak amplitudes and integrals within 40 ms after the stimulus artifact and only if they arose during membrane potential down states. Peak amplitude and integral analysis were performed on each trace, and then presented as a mean of at least 30 whisker-evoked responses. To define up and down states, a membrane potential frequency histogram (1 mV-bin width) was computed for each recorded cell. For each trial, the average membrane potential was determined (10 ms before the stimulus artifact), and if it overlapped with the potentials of the second peak the trace was excluded. Onset latency of PSPs in down state was defined as the time point at which the amplitude exceeded 3 × s.d. of the baseline noise over 5 ms prior to stimulation. It was determined based on an average of at least 20 whisker-evoked PSP traces.

#### Drug application

GABA-A receptors and NMDA receptors were blocked by local and intracellular diffusion of PTX (Sigma, 1 mM) and the NMDA receptor open-channel blocker MK-801 (Tocris, 1 mM) in the recording pipette solution, respectively. The local injection of fluorescent-tag of muscimol was performed as previously described (Gambino et al., 2014). Briefly, mice were anaesthetized with isoflurane and urethane as described above, before being fixed in a stereotaxic frame. Analgesia was provided by local application of lidocaine and i.p. injection of buprenorphine. A burr hole was made to inject the fluorescent muscimol Bodipy(R)-TMR(X) (500 μM in cortex buffer with 5% DMSO, Invitrogen) in the medial part of the posterior thalamic nucleus (POm). The caudal sector of the POm that mainly projects to L1 of S1 (Ohno et al, 2012) was specifically targeted using the following stereotaxic coordinates: RC: −2.00 mm, ML: −1.20 mm, DV: −3.00 mm from the bregma. Glass pipettes (Wiretrol, Drummond) were pulled, back-filled with mineral oil, and front-loaded with the muscimol solution. 100-150 nl were delivered (20 nl/min) using an oil hydraulic manipulator system (MMO-220A, Narishige). For control injection, the same volume of the fluorescent muscimol was injected in thalamic structures that are not involved in somatosensory processing. The craniotomy was then covered with Kwik-Cast (WPI) and mice were prepared for intrinsic optical imaging and whole-cell recordings as described above. To achieve a maximal suppression of neuronal activity, patch-clamp recordings were performed at least one hour after the injection but no longer than 4 hours after the injection. After completion of the experiment, mice were transcardially perfused with 4% paraformaldehyde in PBS (PFA), their brains extracted and post-fixed in PFA overnight. 100-μm coronal brain sections were then made to confirm the site and spread of injections.

### Spatiotemporal analysis of intrinsic optical signal

The intrinsic optical signals were analyzed as previously described (Schubert et al., 2013). The signals were spatially binned (6×6, final resolution: 27.9 μm/pixel or 3×3, final resolution: 13.95 μm/pixel), and a high pass-filter was then applied by subtracting from each image-frame the same image-frame that was convolved using a 1270 μm full-width at half maximum (FWHM) Gaussian kernel. The whisker-evoked intrinsic optical signals were then simulated using a pixel-by-pixel paired t-test, comparing the baseline period and the response period of all trials within a session. The t maps for each individual trial were low pass-filtered with a 340 μm FWHM Gaussian kernel and averaged into a final t map response. A threshold was set to t < −2.0 and any signal below this value was considered to belong to the stimulus-evoked response area. If the pixel value was t ≥ −2.0 it was considered background noise and discarded for quantification. This usually resulted in an image with a clear minimum, representing the response maximum and the barrel’s center of mass. Changes on intrinsic optical signal pixel area caused by whisker trimming were computed as the ratio between the whisker-evoked intrinsic response of the baseline and SWE sessions. All data analysis was performed using a custom software written in MATLAB (MathWorks).

---

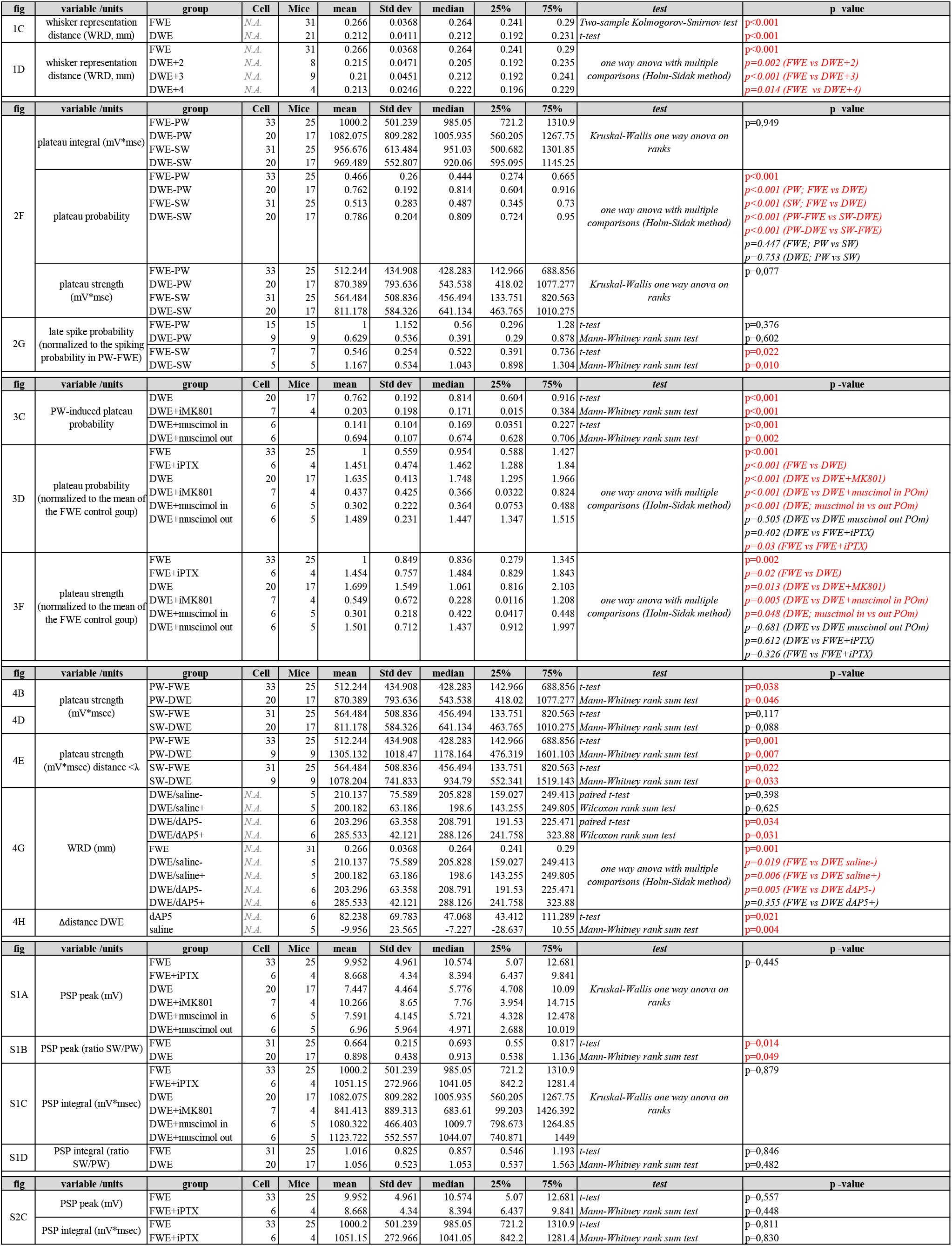

## Notes

### Competing Interest Statement

The authors have declared no competing interest.

## REFERENCES

Alloway, K.D. (2008). Information Processing Streams in Rodent Barrel Cortex: The Differential Functions of Barrel and Septal Circuits. 979–989.

Antic, S.D., Zhou, W.-L., Moore, A.R., Short, S.M., and Ikonomu, K.D. (2010). The decade of the dendritic NMDA spike. J. Neurosci. Res. 88, 2991–3001.

Armstrong-James, M., Welker, E., and Callahan, C.A. (1993). The contribution of NMDA and non-NMDA receptors to fast and slow transmission of sensory information in the rat SI barrel cortex. J. Neurosci. 13, 2149–2160.

Armstrong-James, M., Diamond, M.E., and Ebner, F.F. (1994). An innocuous bias in whisker use in adult rats modifies receptive fields of barrel cortex neurons. J. Neurosci. 14, 6978–6991.

Audette, N.J., Urban-Ciecko, J., Matsushita, M., and Barth, A.L. (2018). POm Thalamocortical Input Drives Layer-Specific Microcircuits in Somatosensory Cortex. Cereb. Cortex 28, 1312–1328.

Brandalise, F., Carta, S., Helmchen, F., Lisman, J., and Gerber, U. (2016). Dendritic NMDA spikes are necessary for timing-dependent associative LTP in CA3 pyramidal cells. Nat. Commun. 7, 13480.

Bureau, I., Saint, F. Von, and Svoboda, K. (2006). Interdigitated Paralemniscal and Lemniscal Pathways in the Mouse Barrel Cortex. 4.

Cardoso, M.M.B., Sirotin, Y.B., Lima, B., Glushenkova, E., and Das, A. (2012). The neuroimaging signal is a linear sum of neurally distinct stimulus- and task-related components. Nat. Neurosci. 15, 1298–1306.

Chen, X., Leischner, U., Rochefort, N.L., Nelken, I., and Konnerth, A. (2011). Functional mapping of single spines in cortical neurons in vivo. Nature 475, 501–505.

Cichon, J., and Gan, W.-B. (2015). Branch-specific dendritic Ca(2+) spikes cause persistent synaptic plasticity. Nature 520, 180–185.

Clem, R.L., and Barth, A. (2006). Pathway-specific trafficking of native AMPARs by in vivo experience. Neuron 49, 663–670.

Clem, R.L., Celikel, T., and Barth, A.L. (2008). Ongoing in Vivo Experience Triggers Synaptic Metaplasticity in the Neocortex. Science (80-.). 319, 101–104.

Deschênes, M., Timofeeva, E., Lavallée, P., and Dufresne, C. (2005). The vibrissal system as a model of thalamic operations. In Progress in Brain Research, pp. 31–40.

Diamond, M.E., Armstrong-James, M., Budway, M.J., and Ebner, F.F. (1992). Somatic sensory responses in the rostral sector of the posterior group (POm) and in the ventral posterior medial nucleus (VPM) of the rat thalamus: Dependence on the barrel field cortex. J. Comp. Neurol. 319, 66–84.

Diamond, M.E., Armstrong-James, M., and Ebner, F.F. (1993a). Experience-dependent plasticity in adult rat barrel cortex. Proc. Natl. Acad. Sci. U. S. A. 90, 2082–2086.

Diamond, M.E., Armstrong-James, M., Ebner, F.F., Armstrong-jamest, M., and Ebner, F.F. (1993b). Experience-dependent plasticity in adult rat barrel cortex. Proc Natl Acad Sci U S A. 90, 2082–2086.

Diamond, M.E., Huang, W., and Ebner, F.F. (1994). Laminar comparison of somatosensory cortical plasticity. Science 265, 1885–1888.

Drew, P.J., and Feldman, D.E. (2009). Intrinsic Signal Imaging of Deprivation-Induced Contraction of Whisker Representations in Rat Somatosensory Cortex. Cereb. Cortex 19, 331–348.

Dringenberg, H.C., Hamze, B., Wilson, A., Speechley, W., and Kuo, M.-C. (2007). Heterosynaptic facilitation of in vivo thalamocortical long-term potentiation in the adult rat visual cortex by acetylcholine. Cereb. Cortex 17, 839–848.

Du, K., Wu, Y.-W., Lindroos, R., Liu, Y., Rózsa, B., Katona, G., Ding, J.B., and Kotaleski, J.H. (2017). Cell-type-specific inhibition of the dendritic plateau potential in striatal spiny projection neurons. Proc. Natl. Acad. Sci. U. S. A. 114.

Feldman, D.E. (2009). Synaptic Mechanisms for Plasticity in Neocortex. Annu. Rev. Neurosci. 32, 33–55.

Feldman, D.E., and Brecht, M. (2005). Map plasticity in somatosensory cortex. Science 310, 810–815.

Feldmeyer, D. (2012). Excitatory neuronal connectivity in the barrel cortex. Front. Neuroanat. 6, 24.

Finnerty, G.T., Roberts, L.S., and Connors, B.W. (1999). Sensory experience modifies the short-term dynamics of neocortical synapses. Nature 400, 367–371.

Fox, K. (2002). Anatomical pathways and molecular mechanisms for plasticity in the barrel cortex. Neuroscience 111, 799–814.

Gainey, M.A., and Feldman, D.E. (2017). Multiple shared mechanisms for homeostatic plasticity in rodent somatosensory and visual cortex. Philos. Trans. R. Soc. B Biol. Sci. 372.

Gambino, F., and Holtmaat, A. (2012). Spike-timing-dependent potentiation of sensory surround in the somatosensory cortex is facilitated by deprivation-mediated disinhibition. Neuron 75, 490–502.

Gambino, F., Pagès, S., Kehayas, V., Baptista, D., Tatti, R., Carleton, A., and Holtmaat, A. (2014). Sensory-evoked LTP driven by dendritic plateau potentials in vivo. Nature 515, 116–119.

Glazewski, S., and Fox, K. (1996). Time course of experience-dependent synaptic potentiation and depression in barrel cortex of adolescent rats. J. Neurophysiol. 75, 1714–1729.

Glazewski, S., Fox, K., Glazewski, S., and Fox, K. (1996). Time Course of Experience-Dependent Synaptic Potentiation and Depression in Barrel Cortex of Adolescent Rats. J. Neurophysiol. 75, 1714–1729.

Glazewski, S., Giese, K.P., Silva, A., and Fox, K. (2000). The role of α-CaMKII autophosphorylation in neocortical experience-dependentplasticity. Nat. Neurosci. 3, 911–918.

Golding, N.L., Staff, N.P., and Spruston, N. (2002). Dendritic spikes as a mechanism for cooperative long-term potentiation. Nature 418, 326–331.

Harding-Forrester, S., and Feldman, D.E. (2018). Somatosensory maps. In Handbook of Clinical Neurology, pp. 73–102.

Holtmaat, A., and Caroni, P. (2016). Functional and structural underpinnings of neuronal assembly formation in learning. Nat. Neurosci. 19, 1553–1562.

House, D.R.C., Elstrott, J., Koh, E., Chung, J., and Feldman, D.E. (2011). Parallel Regulation of Feedforward Inhibition and Excitation during Whisker Map Plasticity. Neuron 72, 819–831.

Jamann, N., Jordan, M., and Engelhardt, M. (2018). Activity-Dependent Axonal Plasticity in Sensory Systems. Neuroscience 368, 268–282.

Jiao, Y., Zhang, C., Yanagawa, Y., and Sun, Q.-Q. (2006). Major effects of sensory experiences on the neocortical inhibitory circuits. J. Neurosci. 26, 8691–8701.

Jones, E.G. (2000). Cortical and subcortical contributions to activity-dependent plasticity in primate somatosensory cortex. Annu. Rev. Neurosci. 23, 1–37.

Jouhanneau, J.-S., Ferrarese, L., Estebanez, L., Audette, N.J., Brecht, M., Barth, A.L., and Poulet, J.F.A. (2014). Cortical fosGFP Expression Reveals Broad Receptive Field Excitatory Neurons Targeted by POm. Neuron 84, 1065–1078.

Keck, T., Scheuss, V., Jacobsen, R.I., Wierenga, C.J., Eysel, U.T., Bonhoeffer, T., and Hübener, M. (2011). Loss of sensory input causes rapid structural changes of inhibitory neurons in adult mouse visual cortex. Neuron 71, 869–882.

Koch, C. (1999). Biophysics of computation: information processing in single neurons (Oxford University Press).

Kock, C.P.J. De, Bruno, R.M., Spors, H., and Sakmann, B. (2009). Layer- and cell-type-specific suprathreshold stimulus representation in rat primary somatosensory cortex. 1, 139–154.

Kubota, Y., Hatada, S., Kondo, S., Karube, F., and Kawaguchi, Y. (2007). Neocortical inhibitory terminals innervate dendritic spines targeted by thalamocortical afferents. J. Neurosci. 27, 1139–1150.

Larkum, M. (2013). A cellular mechanism for cortical associations: an organizing principle for the cerebral cortex. Trends Neurosci. 36, 141–151.

Larkum, M.E., Zhu, J.J., and Sakmann, B. (1999). A new cellular mechanism for coupling inputs arriving at different cortical layers. Nature 398, 338–341.

Larkum, M.E., Waters, J., Sakmann, B., and Helmchen, F. (2007). Dendritic Spikes in Apical Dendrites of Neocortical Layer 2/3 Pyramidal Neurons. J. Neurosci. 27, 8999–9008.

Larkum, M.E., Nevian, T., Sandler, M., Polsky, A., and Schiller, J. (2009). Synaptic Integration in Tuft Dendrites of Layer 5 Pyramidal Neurons: A New Unifying Principle. Science (80-.). 325, 756–760.

Lavallée, P., Urbain, N., Dufresne, C., Bokor, H., Acsády, L., and Deschênes, M. (2005). Feedforward inhibitory control of sensory information in higher-order thalamic nuclei. J. Neurosci. 25, 7489–7498.

Li, L., Gainey, M.A., Goldbeck, J.E., and Feldman, D.E. (2014). Rapid homeostasis by disinhibition during whisker map plasticity. Proc. Natl. Acad. Sci. 111, 1616–1621.

Major, G., Polsky, A., Denk, W., Schiller, J., and Tank, D.W. (2008). Spatiotemporally Graded NMDA Spike/Plateau Potentials in Basal Dendrites of Neocortical Pyramidal Neurons. J. Neurophysiol. 99, 2584–2601.

Masri, R., Trageser, J.C., Bezdudnaya, T., Li, Y., and Keller, A. (2006). Cholinergic regulation of the posterior medial thalamic nucleus. J. Neurophysiol. 96, 2265–2273.

Masri, R., Bezdudnaya, T., Trageser, J.C., and Keller, A. (2008). Encoding of stimulus frequency and sensor motion in the posterior medial thalamic nucleus. J. Neurophysiol. 100, 681–689.

Masri, R., Quiton, R.L., Lucas, J.M., Murray, P.D., Thompson, S.M., and Keller, A. (2009). Zona incerta: a role in central pain. J. Neurophysiol. 102, 181–191.

Mease, R.A., Metz, M., and Groh, A. (2016). Cortical Sensory Responses Are Enhanced by the Higher-Order Thalamus. Cell Rep. 14, 208–215.

Oberlaender, M., Ramirez, A., and Bruno, R.M. (2012). Sensory experience restructures thalamocortical axons during adulthood. Neuron 74, 648–655.

Ohno, S., Kuramoto, E., Furuta, T., Hioki, H., Tanaka, Y.R., Fujiyama, F., Sonomura, T., Uemura, M., Sugiyama, K., and Kaneko, T. (2012). A morphological analysis of thalamocortical axon fibers of rat posterior thalamic nuclei: a single neuron tracing study with viral vectors. Cereb. Cortex 22, 2840–2857.

Palmer, L., Murayama, M., and Larkum, M. (2012). Inhibitory Regulation of Dendritic Activity in vivo. Front. Neural Circuits 6, 26.

Palmer, L.M., Shai, A.S., Reeve, J.E., Anderson, H.L., Paulsen, O., and Larkum, M.E. (2014). NMDA spikes enhance action potential generation during sensory input.

Petersen, C.C.H., Grinvald, A., and Sakmann, B. (2003). Spatiotemporal dynamics of sensory responses in layer 2/3 of rat barrel cortex measured in vivo by voltage-sensitive dye imaging combined with whole-cell voltage recordings and neuron reconstructions. J. Neurosci. 23, 1298–1309.

Petreanu, L., Mao, T., Sternson, S.M., and Svoboda, K. (2009). The subcellular organization of neocortical excitatory connections. Nature 457, 1142–1145.

Polley, D.B., Chen-Bee, C.H., and Frostig, R.D. (1999). Two directions of plasticity in the sensory-deprived adult cortex. Neuron 24, 623–637.

Salt, T.E. (1986). Mediation of thalamic sensory input by both NMDA receptors and non-NMDA receptors. Nature 322, 263–265.

Schneider, C.A., Rasband, W.S., and Eliceiri, K.W. (2012). NIH Image to ImageJ: 25 years of image analysis. Nat. Methods 9, 671–675.

Schubert, V., Lebrecht, D., and Holtmaat, A. (2013). Peripheral deafferentation-driven functional somatosensory map shifts are associated with local, not large-scale dendritic structural plasticity. J. Neurosci. 33, 9474–9487.

Sermet, B.S., Truschow, P., Feyerabend, M., Mayrhofer, J.M., Oram, T.B., Yizhar, O., Staiger, J.F., and Petersen, C.C.H. (2019). Pathway-, layer-and cell-type-specific thalamic input to mouse barrel cortex. Elife 8.

Sherman, S.M. (2017). Functioning of circuits connecting thalamus and cortex. Compr. Physiol. 7, 713–739.

Stern, E. a, Maravall, M., and Svoboda, K. (2001). Rapid development and plasticity of layer 2/3 maps in rat barrel cortex in vivo. Neuron 31, 305–315.

Takahashi, N., Oertner, T.G., Hegemann, P., and Larkum, M.E. (2016). Active cortical dendrites modulate perception. Science (80-.). 354, 1587–1590.

Trageser, J.C., and Keller, A. (2004). Reducing the uncertainty: gating of peripheral inputs by zona incerta. J. Neurosci. 24, 8911–8915.

Trageser, J.C., Burke, K.A., Masri, R., Li, Y., Sellers, L., and Keller, A. (2006). State-dependent gating of sensory inputs by zona incerta. J. Neurophysiol. 96, 1456–1463.

Ts’o, D.Y., Frostig, R.D., Lieke, E.E., and Grinvald, A. (1990). Functional organization of primate visual cortex revealed by high resolution optical imaging. Science 249, 417–420.

van Versendaal, D., Rajendran, R., Saiepour, M.H., Klooster, J., Smit-Rigter, L., Sommeijer, J.-P., De Zeeuw, C.I., Hofer, S.B., Heimel, J.A., and Levelt, C.N. (2012). Elimination of Inhibitory Synapses Is a Major Component of Adult Ocular Dominance Plasticity. Neuron 74, 374–383.

Veinante, P., and Deschênes, M. (1999). Single- and multi-whisker channels in the ascending projections from the principal trigeminal nucleus in the rat. J. Neurosci. 19, 5085–5095.

Viaene, A.N., Petrof, I., and Sherman, S.M. (2011). Properties of the thalamic projection from the posterior medial nucleus to primary and secondary somatosensory cortices in the mouse. Proc. Natl. Acad. Sci. U. S. A. 108, 18156–18161.

Vincis, R., Lagier, S., Van De Ville, D., Rodriguez, I., and Carleton, A. (2015). Sensory-Evoked Intrinsic Imaging Signals in the Olfactory Bulb Are Independent of Neurovascular Coupling. Cell Rep. 12, 313–325.

Wallace, D.J., and Sakmann, B. (2008). Plasticity of representational maps in somatosensory cortex observed by in vivo voltage-sensitive dye imaging. Cereb. Cortex 18, 1361–1373.

Wilent, W.B., and Contreras, D. (2004). Synaptic responses to whisker deflections in rat barrel cortex as a function of cortical layer and stimulus intensity. J. Neurosci. 24, 3985–3998.

Williams, L.E., and Holtmaat, A. (2019). Higher-Order Thalamocortical Inputs Gate Synaptic Long-Term Potentiation via Disinhibition. Neuron 101, 91–102.e4.

Williams, S.R., and Fletcher, L.N. (2019). A Dendritic Substrate for the Cholinergic Control of Neocortical Output Neurons. Neuron 101, 486–499.e4.

Wimmer, V.C., Broser, P.J., Kuner, T., and Bruno, R.M. (2010a). Experience-induced plasticity of thalamocortical axons in both juveniles and adults. J. Comp. Neurol. 518, 4629–4648.

Wimmer, V.C., Bruno, R.M., de Kock, C.P.J., Kuner, T., and Sakmann, B. (2010b). Dimensions of a projection column and architecture of VPM and POm axons in rat vibrissal cortex. Cereb. Cortex 20, 2265–2276.

Wolfe, J., Houweling, A.R., and Brecht, M. (2010). Sparse and powerful cortical spikes. Curr. Opin. Neurobiol. 20, 306–312.

Xu, N., Harnett, M.T., Williams, S.R., Huber, D., O’connor, D.H., Svoboda, K., and Magee, J.C. (2012). Nonlinear dendritic integration of sensory and motor input during an active sensing task. Nature 492, 247–251.

Yazaki-Sugiyama, Y., Kang, S., Cteau, H., Fukai, T., and Hensch, T.K. (2009). Bidirectional plasticity in fast-spiking GABA circuits by visual experience. Nature 462, 218–221.

Yu, X., Chung, S., Chen, D.-Y., Wang, S., Dodd, S.J., Walters, J.R., Isaac, J.T.R., and Koretsky, A.P. (2012). Thalamocortical Inputs Show Post-Critical-Period Plasticity. Neuron 74, 731–742.

Zhang, W., and Bruno, R.M. (2019). High-order thalamic inputs to primary somatosensory cortex are stronger and longer lasting than cortical inputs. Elife 8.

